# Acute thermal stress elicits interactions between gene expression and alternative splicing in a fish of conservation concern

**DOI:** 10.1101/2022.02.20.481219

**Authors:** Matt J Thorstensen, Andy J Turko, Daniel D Heath, Ken M Jeffries, Trevor E Pitcher

## Abstract

Transcriptomics provides a mechanistic understanding of an organism’s response to environmental challenges such as increasing temperatures, which can provide key insights into the threats posed by thermal challenges associated with urbanization and climate change. Differential gene expression and alternative splicing are two elements of the transcriptomic stress response that may work in tandem, but relatively few studies have investigated these interactions in fishes of conservation concern. We studied the imperilled redside dace (*Clinostomus elongatus*) as thermal stress is hypothesised to be an important cause of population declines. We tested the hypothesis that gene expression-splicing interactions contribute to the thermal stress response. Wild fish exposed to acute thermal stress were compared with both handling controls and fish sampled directly from a river. Liver tissue was sampled to study the transcriptomic stress response. Thermally stressed fish showed a prominent transcriptional response (estimated with mRNA transcript abundance) related to transcription regulation and responses to unfolded proteins, and prominent alternatively spliced genes related to gene expression regulation and metabolism. One splicing factor, *prpf38b*, was upregulated in the thermally stressed group compared to the other treatments. This splicing factor may have a role in the Jun/AP-1 cellular stress response, a pathway with wide-ranging and context-dependent effects. Given large gene interaction networks and the context-dependent nature of transcriptional responses, our results highlight the importance of understanding interactions between gene expression and splicing for understanding transcriptomic responses to thermal stress. Our results also reveal transcriptional pathways that can inform conservation breeding, translocation, and reintroduction programs for redside dace and other imperilled species by identifying appropriate source populations.

**SUMMARY STATEMENT:** Gene expression and alternative splicing interact in response to thermal stress in an imperilled fish, with implications for conservation and mechanisms of thermal tolerance in vertebrate ectotherms.

## INTRODUCTION

Environmental temperature influences many aspects of the physiology and behaviour of ectothermic animals (Fry, 1947; Schulte, 2015). The thermal environment, especially maximum temperatures, is therefore one of the most important factors that determines the fundamental niche, and thus geographic distribution, of many ectotherms (Bennett et al., 2021; Bozinovic et al., 2011; Day et al., 2018). Aquatic systems are especially vulnerable to increasing temperatures in conjunction with other factors, which can threaten resource availability and biodiversity (Dudgeon, 2019). Anthropogenic disturbances, ranging in scope from global climate change to local land use changes, have increased maximum water temperatures in many aquatic systems, and these temperature extremes are predicted to become more severe (O’Reilly et al., 2015). This warming is hypothesised to be a threat to the distribution and even long-term persistence of many aquatic species (Heino et al., 2009; Myers et al., 2017). However, there is a high degree of interspecific variation in thermal sensitivity even among species that share a common habitat, and the underlying physiological mechanisms are poorly understood (Komoroske et al., 2021; Pörtner et al., 2017). An improved mechanistic understanding of responses to high temperatures is important for predicting population responses to thermal challenges and for guiding recovery actions, particularly for species at risk (e.g., Eliason et al., 2011; Lefevre et al., 2021; McDonnell et al., 2021; Wenger et al., 2011).

Transcriptomics has emerged as a powerful tool for characterizing the mechanisms of organismal response to stressors such as high temperature, which can then be applied to conservation management (Connon et al., 2018). Comparisons of molecular responses among populations can reveal the mechanisms underlying vulnerable and resistant populations to a stressor, with implications for managing habitat and guiding reintroduction (e.g., Jeffries et al., 2019). While mRNA abundance most accurately reflects gene transcription (Buccitelli and Selbach, 2020; Jeffries et al., 2021), tests of differential abundance of mRNA transcripts are commonly referred to as tests of differential gene expression in transcriptomics studies (e.g., Conesa et al., 2016). In addition to tests of mRNA abundance, RNA-sequencing also enables tests of alternative splicing (Salisbury et al., 2021). Instead of mRNA abundance changing in response to a stressor as in transcriptomics, exons within genes are differentially assembled in post-transcriptional modifications of RNA.

While mRNA abundance is better understood than alternative splicing in the eukaryotic stress response (Salisbury et al., 2021), splicing also contributes to controlling stress responses in plants, yeast, fruit flies, shrimp, and humans (Chaudhary et al., 2019; De Nadal et al., 2011; Kornblihtt et al., 2013; Laloum et al., 2018; Zhang et al., 2019). Furthermore, recent studies indicate that alternative splicing is a key element of the response to environmental change in fishes, such as salinity changes (Thorstensen et al., 2021), acute hypoxia (Xia et al., 2018), cold acclimation (Healy and Schulte, 2019), cold stress (Li et al., 2020), and heat stress (Tan et al., 2019). Differential splicing also contributes to evolutionary change including local adaptation of ecotypes (Jacobs and Elmer, 2021; Salisbury et al., 2021) and speciation (Singh et al., 2017; Terai et al., 2003). However, relatively little is known about changes in splicing following an acute thermal stress event (Tan et al., 2019).

Gene expression and alternative splicing have often been studied separately, but recent work suggests these mechanisms should be considered in tandem (e.g., Healy and Schulte, 2019; Jacobs and Elmer, 2021; Singh and Ahi, 2022). This combined approach can reveal interactions between splicing and gene expression, such as by differentially expressed splicing factors that contribute to downstream splicing. For instance, transcription cofactor binding genes were alternatively spliced, while genes involved in the spliceosome were differentially expressed in a cold stress experiment in Nile tilapia (*Oreochromis niloticus*) (Li et al., 2020). Different splicing factors have been found to affect gene expression in contexts outside of the thermal stress response, sometimes referred to as cross-talk between gene expression and the spliceosome (Änkö, 2014; Kim et al., 2018; Smith et al., 1989). Therefore, we hypothesized that gene expression-splicing interactions may also contribute to the acute heat stress response.

To understand the transcriptome-level interactions between differential gene expression and alternative splicing, we studied the regionally imperilled redside dace (*Clinostomus elongatus*). This cyprinid inhabits cool-water streams in northeastern North America, but population sizes and range areas have declined dramatically (COSEWIC, 2017). Redside dace are considered endangered in Canada (COSEWIC, 2017; Redside Dace Recovery Team, 2010) and many populations are considered imperilled in the United States (Serrao et al., 2018). Several studies suggest that redside dace population declines may be linked to thermal stress resulting from the combined effects of urbanization and climate change, and that thermal tolerance varies among genetically distinct redside dace populations (Leclair et al., 2020; Turko et al., 2020; Turko et al., 2021). Understanding the mechanistic basis of these differences is important for predicting thermal responses for different populations and for guiding potential conservation programs such as translocation or reintroduction based on captively bred individuals. Thus, in addition to our main goal of testing the overarching “expression-splicing interaction” hypothesis, we also aimed to identify thermally responsive genes of redside dace that can be applied to future conservation reintroduction programs for this imperilled species.

Using individuals directly from their natal stream, we experimentally investigated the molecular mechanisms underlying acute thermal stress in a wild population of redside dace (see Figure 1). We sampled livers from adult redside dace exposed to acute thermal stress (following a standard critical thermal maximum (CTmax) protocol; Turko et al., 2020) and two control groups: “wild”, fish sampled immediately after capture from the stream, and “handling control”, fish that were treated the same as thermally stressed fish but kept at ambient temperatures. We used RNA sequencing to profile differentially expressed and alternatively spliced genes unique to thermal stress to understand the molecular mechanisms that redside dace use to respond to acute thermal stress. Differential gene expression was tested with mRNA abundance, while alternative splicing was assessed with differentially used exons from within mRNA transcripts. Our hypothesis was that the thermal response involves interactions between splicing and gene expression. This hypothesis predicts that splicing factors that show differential expression in response to a thermal challenge are among the specific genes that enable interactions between splicing and gene expression. Therefore, splicing factors upregulated in the thermally stressed fish compared to both other groups were analyzed for possible connections with stress and thermal response genes.

**Figure 1.**
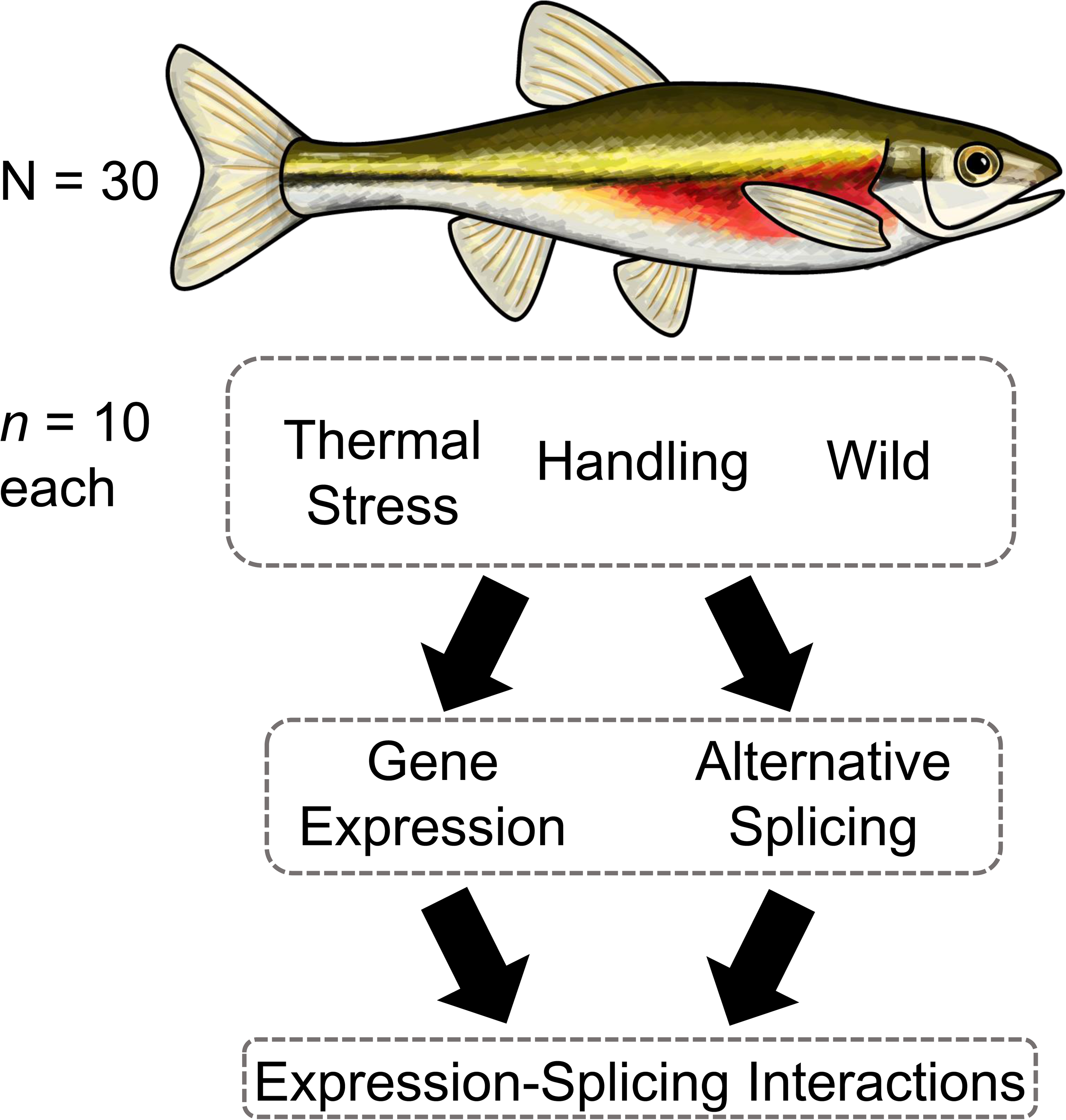
Conceptual diagram of experimental design and analysis approaches. Redside dace (*Clinostomus elongatus*) were divided into three experimental treatments of *n*=10 individuals each: CTmax as a thermal stressor, a handling treatment where fish were handled as in the CTmax protocol but not heated, and a wild control. Messenger RNA sequencing was performed on liver tissue of all individuals. Genes showing differential expression and alternative splicing were analyzed, with particular emphasis on the thermal stress treatment compared to both others. We hypothesized that gene expression-splicing interactions may contribute to the thermal stress response, and analyzed differentially expressed splicing factors in the thermal stress treatment. Spliced genes associated with the splicing factors were also analyzed, with implications for expression-splicing interactions in the context of thermal stress.

## METHODS

### Fieldwork and thermal stress

Adult redside dace (N=30) were collected using a seine net from a single pool in the Kokosing River, Ohio, USA, (40°32’43.1”N 82°39’15.2”W) over two days in February 2019. Fish were then randomly assigned to one of three treatments (each *n* = 10): “wild” fish, thermal stress, or handling control (Figure 1). Wild fish were euthanized via blunt force trauma to the head and spinal severance within 1 min of capture, the body cavity was opened with a ventral incision, and fish were submerged in a high salt solution (700 g/L ammonium sulfate, 25 mM sodium citrate, 20 mM ethylenediaminetetraacetic acid, pH 5.2; Wellband and Heath, 2017) to preserve tissues for transcriptomic analysis. Tissues were stored first at 4°C for 48h to facilitate preservation, and were subsequently stored at -20°C until RNA extraction. For the thermally stressed group, fish were subjected to a standard critical thermal maximum protocol as described in detail elsewhere (Turko et al., 2020). Briefly, fish were quickly (within 10 min of capture) transferred to individual mesh-walled plastic containers submerged in an aerated, thermostatically controlled water bath (VWR model 1203) filled with river water. After a 15 min acclimation period, water temperature was raised by 0.33°C/min until fish could not maintain equilibrium (upright position) in the water column for 3s. Temperature and dissolved oxygen (always >80%) were monitored throughout each experiment (YSI Pro Plus multi-parameter instrument, Yellow Springs, OH, USA). Once fish lost equilibrium (75-85 min), the temperature was recorded and fish were immediately euthanized and preserved as described above. Fish in the handling control group were treated identically to thermally stressed fish except they did not experience increased water temperatures. Instead, these fish were sampled after the average length of a thermal stress experiment (∼80 min). Hereafter, fish in the wild group are referred to as “wild”, fish that underwent CTmax as the “thermal stress” group, and fish that were in the handling control as “handled”.

### RNA extraction and sequencing

RNA was extracted from fish liver using RNeasy Plus Mini Prep Kits (QIAGEN) following manufacturer protocols. Total RNA was sent to the Génome Québec Innovation Centre sequencing facility (http://gqinnovationcenter.com), where 250 nanograms of total RNA per fish were used with the NEBNext Poly(A) Magnetic Isolation Module (New England BioLabs). RNA integrity number (RIN) scores assessed with a Bioanalyzer (Agilent) were >7 for all fish (8.83 mean, ±0.67 standard deviation). Stranded cDNA libraries were created with the NEBNext Ultra II Directional RNA Library Prep Kit for Illumina (New England Biolabs). The 30 fish were sequenced for 100 base pair reads on one lane of a NovaSeq 6000 (Illumina). A mean of 50.5 million reads per sample were sequenced (±9.2 million standard deviation) (Supplementary Table S1).

### Transcriptome assembly and annotation

Raw reads were trimmed with Trimmomatic version 0.36, where reads under 36 base pairs long were discarded, leading and trailing base pairs were discarded with Phred scores lower than 5, and a sliding window was of four base pairs was used where the window was removed if the average read quality fell below five (Bolger et al., 2014). Read quality metrics before and after trimming were checked with FastQC version 0.11.8 and multiQC version 1.9 (Andrews, 2010; Ewels et al., 2016). Following trimming, a mean 49.9 million reads per sample were retained (±9.2 million s.d.) (Supplementary Table S1). Trinity version 2.9 was used to assemble the transcriptome with trimmed reads with default options, followed by BUSCO version 3.0.2 with the ray-finned fish lineage (actinopterygii_odb10) to assess transcriptome completeness (Grabherr et al., 2011; Seppey et al., 2019). Trinotate version 3.2.0 was used for transcriptome annotation following software guidelines, except RNAMMER was not used with these data (https://github.com/Trinotate/Trinotate.github.io/wiki/Software-installation-and-data-required) (Bryant et al., 2017). In short, the NCBI blastx and blastp databases were used to search transcripts and predicted proteins, respectively, HMMER version 3.3 was used to identify protein families, signalP version 4.1 was used to identify signal peptides, and tmhmm version 2 was used to identify transmembrane helices (Altschul et al., 1990; Krogh et al., 2001; Petersen et al., 2011; Wheeler and Eddy, 2013) . All results were collected in Trinotate for transcriptome annotations. Following Pearson (2013), annotations were filtered for those with E-values <1 x 10^-6^ and bit scores >50.

The SuperTranscripts pipeline was used because of its potential for describing differential exon usage; SuperTranscript results were thus also used for gene expression (Davidson et al., 2017). Here, Salmon version 1.1.0 with the --dumpEq option was used for initial transcript quantification against the reference transcriptome (Patro et al., 2017). Equivalence classes from Salmon were used in Corset version 1.07 to generate super-clusters with five nucleotides minimum required to overlap between transcripts, and super-clusters were discarded if over 1000 contigs aligned to them (Davidson and Oshlack, 2014). A linear representation of the transcriptome was generated with the Corset outputs and the Trinity transcriptome using Lace version 1.14.1 (https://github.com/Oshlack/Lace).

### Differential Gene Expression

Corset counts for super-clusters were used for differential gene expression (DGE) using edgeR version 3.30.3 (Robinson et al., 2010). While the counts from Corset super-clusters reflect gene transcription (Buccitelli and Selbach, 2020; Jeffries et al., 2021), the resulting statistical tests are referred to here as DGE in a manner consistent with similar literature (e.g., Conesa et al., 2016). Count data were first filtered for genes showing any expression. Then, a genewise negative binomial generalized linear model with quasi-likelihood test (glmQLFit) was used after data normalization and robust dispersion estimation. The design formula for the model included experimental group (wild, handled, thermally stressed) and RNA integrity number to explicitly model differences in RNA quality between samples (Supplementary Table S1) (Gallego Romero et al., 2014). Pairwise comparisons were drawn between each experimental treatment using genewise negative binomial generalized linear models with quasi-likelihood tests (glmQLFtest). Only clusters significant at a Benjamini-Hochberg adjusted false discovery rate (*q*)<0.05 were retained for downstream functional analyses (Benjamini and Hochberg, 1995). In addition, clusters with higher or lower expression in the thermal stress treatment compared to both the wild and handled treatments (i.e., |log_2_-fold change|>0 compared to both wild and handled) were retained as exhibiting ‘thermal stress-specific’ expression. Multidimensional scaling as implemented in edgeR and a heatmap were used to visualize broad patterns of differential gene expression among all clusters and those specific to thermal stress, respectively.

To find summary gene ontology (GO) terms represented by differentially expressed and spliced super-clusters, we used the EnrichR version 2.1 databases Biological Process 2018, Molecular Function 2018, and Cellular Component 2018 (Kuleshov et al., 2016). GO terms were analyzed in pairwise comparisons between each experimental treatment in both the DGE and DEU results. Because we were interested in patterns of splicing and expression with respect to thermal tolerance, results unique to the thermal stress experimental treatment were given special attention. For DGE, statistically significant clusters (*q*<0.05) that were either upregulated (log_2_-fold changes>0) or downregulated (log_2_-fold changes<0) in the thermally stressed treatment with respect to both the handled control and wild group were retained for these thermal stress-specific results, in addition to overall thermal stress-specific genes (|log_2_-fold change|>0). Gene set enrichment analysis was conducted with overall thermal stress-specific genes, because the upregulation of certain genes may downregulate given pathways, and downregulation of other genes may upregulate pathways (Reynolds et al., 2013). For visualization, non-redundant GO terms for genes that showed thermal stress-specific expression were explored with Revigo where significant GO terms and adjusted *p*-values were used with an allowed semantic similarity of 0.7, and terms were searched against the whole UniProt database (Supek et al., 2011).

### Early Response Genes

To investigate the possibility that thermally stressed or handed fish exhibited gene expression changes indicative of an acute stress response, several early response genes were explored in the DGE data. These were clusters annotated to the genes transcription factor Jun/AP-1 (*jun*), transcription factor jun-B (*jun-B*), transcription factor jun-D (*jun-D*), immediate early response gene 2 (*ier2*), myc proto-oncogene (*myc*), proto-oncogene c-Fos (*c-Fos*), and metallothiol transferase FosB (*fosB*). The panel of early response genes represents a positive control of genes we expected would change in expression if the thermal and handling stressors were reflected in a transcriptomic response (Bahrami and Drabløs, 2016; Fowler et al., 2011; Jeffries et al., 2018; Sopinka et al., 2016). Therefore, if differential expression was observed in these genes, other genes that were differentially expressed between the handled or thermally stressed treatments can be assumed to be related to the specific stressors of each condition.

### Differential Exon Usage

Differential exon usage (DEU), or the relative usage of exons within genes, was estimated using STAR version 2.7.3a to create splice junction files for each individual (SJ.out.tab), which were concatenated into one splice junction file (Dobin et al., 2013). STAR was run in two pass mode with all reads mapped on the first pass. The Mobius.py script in Lace version 1.14.1 was used to create a .gtf file from the Lace-clustered transcriptome and the splice junction file from STAR. Then, the featureCounts function in Subread version 2.01 was used with fractional counts (--fraction), where input files were the new splice junction-specific .gtf file, the super-clusters count file from Corset, and the aligned .bam files from STAR to generate exon counts (Liao et al., 2013). DEU was tested for with DEXseq version 1.34.1 (Anders et al., 2012). As with tests for DGE, RIN scores were used but with a centered and scaled mean around 0 for generalized linear model convergence. The design formula for DEU included the individual fish, scaled RIN, exon expression, and experimental treatment in interaction with exon expression. After estimating size factors and dispersions, exon usage coefficients were estimated by being fit to experimental treatments. Only exons with differential expression significant at an *q*<0.05 were retained for downstream function analyses. Similar to DGE analyses, exons with higher or lower expression in the thermal stress treatment compared to both the wild and handled treatments (i.e., |log_2_-fold change|>0 compared to both wild and handled) were retained as exhibiting ‘thermal stress-unique’ expression. These steps were performed following guidelines in the Lace GitHub repository (https://github.com/Oshlack/Lace/wiki/Example:-Differential-Transcript-Usage-on-a-non-model-organism).

Only GO terms from the Biological Process 2018, Molecular Function 2018, and Cellular Component 2018 databases with *q*<0.05 were retained for further analyses. As with GO terms represented by DGE, Revigo was used to explore non-redundant GO terms for exons that showed thermal stress-specific expression (Supek et al., 2011).

### Gene Expression-Splicing Interactions

To explicitly evaluate our hypothesis that alternative splicing is an important mechanism used by redside dace responding to a thermal challenge, we focused on splicing factors uniquely upregulated (DGE log_2_-fold changes>0 when compared to both other groups) in fish in the thermal stress treatment. Using the STRING version 11.0 database (Szklarczyk et al., 2019), we analyzed genes in molecular pathways with the splicing factors identified previously using the *Danio rerio* database. Here, genes with significant DEU were identified as possibly important for thermal stress response.

## RESULTS

### Transcriptome assembly and annotation

Trinity assembled unaligned reads into a transcriptome of 714,933 unique transcripts in 429,016 unique genes with a BUSCO score for transcriptome completeness of 89.8%. Of these putative transcripts and genes, 155,547 transcripts representing 59,755 genes were annotated using Trinotate and associated programs after filtering for E-values <1 x 10^-6^ and bit scores >50. Corset clustered transcripts from Trinity into 83,217 super-clusters representing 143,841 clusters.

### Differential Gene Expression

Of the 143,841 clusters from Corset irrespective of available annotations, 46,140 had measurable expression in any single individual. Between the thermal stress group and handled control, 1,531 clusters showed significant DGE, 786 with relatively higher expression in thermal stress and 745 with relatively lower expression in thermal stress compared with the handled control (Table 1; Supplementary Table S2). Between the thermal stress and wild group, 6,770 clusters showed significant DGE, 3,992 with relatively higher expression in thermal stress and 2,778 with lower expression in thermal stress compared with the wild group (Supplementary Table S3). For clusters with expression unique to thermal stress, 579 showed positive DGE compared to the two other groups, while 559 showed negative DGE compared to the two other groups (Supplementary Table S4; Figure 2). Multidimensional scaling with all clusters and a heatmap of counts per million for each of 1,138 clusters showing significant differential gene expression unique to the thermal stress treatment (579 positive, 559 negative; *q* < 0.05) reveal a gradient in expression response from the wild group to the handled control, and the thermal stress group (Figure 2). While the fish ‘wild 3’ was an outlier in mRNA abundance profile and had the lowest RIN score out of all individuals of 7.1 (Figure 2; Supplementary Table 1), its removal from differential gene expression analyses did not qualitatively affect downstream results. Rather than introduce bias by removing this outlier individual, it was retained for all analyses.

**Figure 2.**
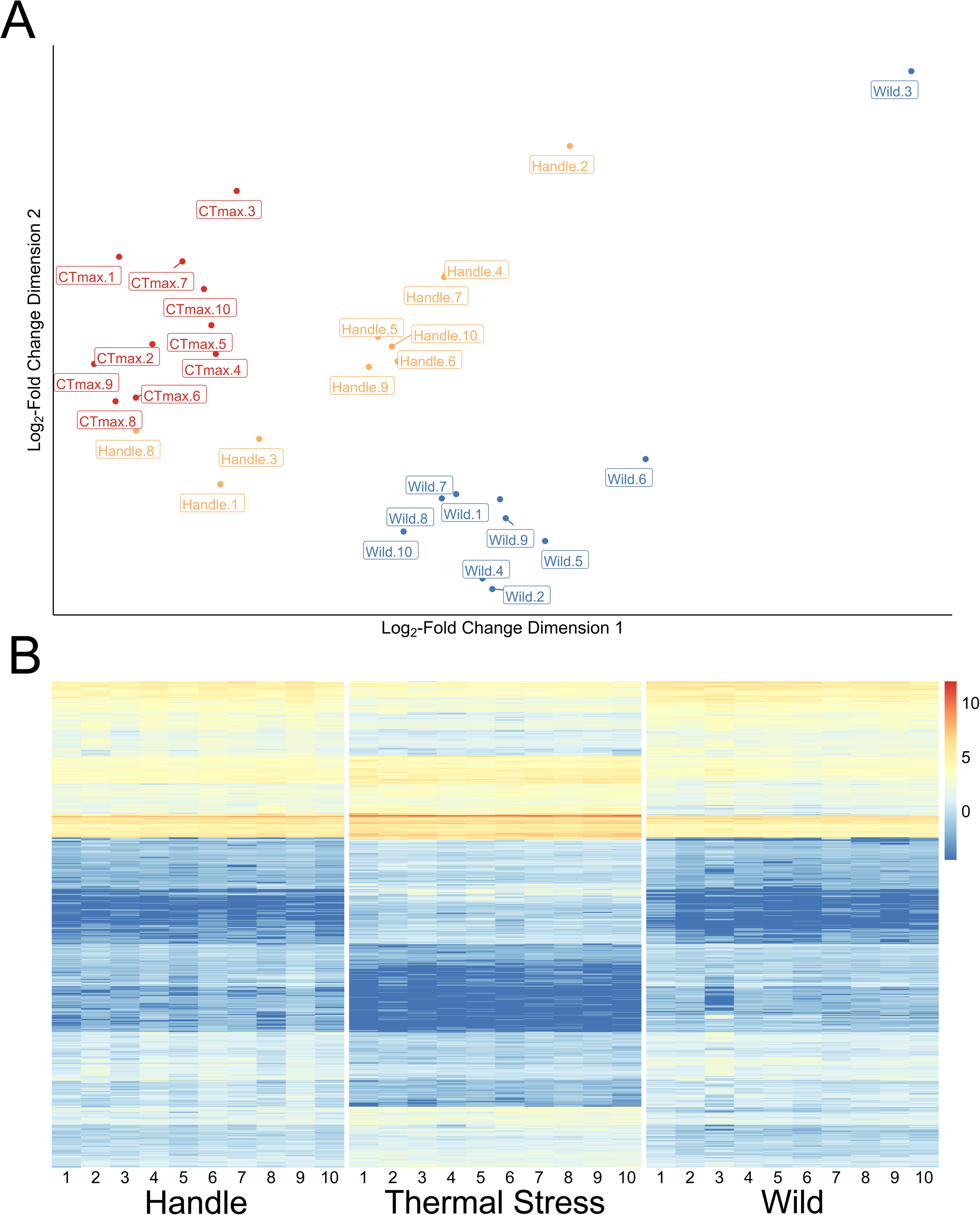
Differential gene expression in response to thermal stress. (A) Visualization of cluster expression as implemented by a multidimensional scaling (MDS) plot using edgeR, where distances between plots are approximations of log_2_-fold changes between samples. Input data are cluster (∼transcript) expression counts filtered for any expression among any of the *n*=30 individuals in the experiment. Individual labels are comprised of the experimental treatment or control (thermally stressed (abbreviated as CTmax), handled, or wild) and the individual’s identifying number. (B) Heatmap of counts per million for each of 1,138 clusters showing significant differential gene expression unique to the thermal stress treatment (579 positive, 559 negative; *q* < 0.05). That is, each cluster included in this plot either shows higher expression in the thermal stress treatment compared to both the wild and handled controls, or lower expression in the thermal stress treatment compared to both controls. Individuals are groups by experimental treatment, and numbers identifying individuals within each treatment are on the X-axis.

**Table 1.** Summary table of pairwise results for differential gene expression among three experimental treatments. Clusters (∼transcripts) were identified and quantified with Corset, and differential gene expression (DGE) was analyzed with edgeR. EnrichR was used to summarize annotated clusters under different pairwise comparisons into gene ontology (GO) terms, among three databases: Biological Process 2018, Molecular Function 2018, and Cellular Component 2018. Counts of clusters associated to known genes are reported as Genes. Positive and negative expression for clusters and GO terms are relative to the pairwise comparison used; positive expression represents clusters higher in the first treatment of a comparison, while negative expression represents clusters higher in the second treatment of a comparison. The thermal stress treatment is abbreviated as CTmax.

|  | **Wild vs Handled** | **Wild vs CTmax** | **CTmax vs Handled** |
| --- | --- | --- | --- |
| **Number of Significant DGE Clusters Overall** | 2362 | 6770 | 1531 |
| **Number of Significant Positive DGE Clusters** | 578 | 2778 | 786 |
| **Number of Significant Negative DGE Clusters** | 1784 | 3992 | 745 |
| **Positive DGE Genes** | 263 | 1478 | 328 |
| **Negative DGE Genes** | 670 | 1682 | 218 |
| **Positive DGE Biological Process GO Terms** | 0 | 98 | 27 |
| **Positive DGE Molecular Component GO Terms** | 0 | 5 | 12 |
| **Positive DGE Cellular**<br>**Component GO Terms** | 0 | 40 | 0 |
| **Negative DGE Biological Process**<br>**GO Terms** | 60 | 211 | 0 |
| **Negative DGE Molecular**<br>**Component GO Terms** | 32 | 43 | 0 |
| **Negative DGE Cellular**<br>**Component GO Terms** | 1 | 1 | 0 |

Among annotated clusters showing significant DGE, 328 genes were identified as showing relatively higher expression in thermal stress compared to the handled control, and 218 genes lower for the thermal stress group (Table 1; Supplementary Table S2). Between the thermal stress treatment and handled control, 39 GO terms were identified from genes showing higher expression for thermal stress (no GO terms were found for genes with higher expression in handled control) (Supplementary Table S7). Between the thermal stress and wild groups of fish, 1,682 genes were higher for thermal stress (1,478 higher for wild group) (Table 1; Supplementary Table S3). Between the thermal stress treatment and wild group 256 GO terms were identified from genes showing higher expression in thermal stress, while 143 genes had higher expression in wild fish (Supplementary Table S8).

For genes that showed thermal stress-specific expression (i.e., |log_2_-fold changes| > 0 compared to both other groups for thermal stress-specific expression, respectively), 579 were identified as showing higher expression in thermal stress compared to both controls (216 annotated clusters), and 559 showed lower expression in thermal stress compared to both controls (103 annotated clusters) (Supplementary Table S4). For GO terms related to genes specific to thermal stress, 32 GO terms were identified among genes with positive expression (21 Biological Process terms, 11 Molecular Function terms) while no GO terms were identified for genes with negative expression (Supplementary Figure S1; Supplementary Table S9). Using Revigo with the thermal stress-specific GO terms, 12 Biological Process GO terms and 8 Molecular Function GO terms were retained for visualization (Supplementary Figure S1). With the 1,138 total clusters identified as unique to thermal stress (775 annotated), 37 Biological Process, 30 Molecular Function, and one Cellular Component GO terms were identified. With Revigo, 25 Biological Process, 21 Molecular Function, and one Cellular Component non-redundant GO terms were retained for visualization (Figure 3). The GO terms regulation of transcription, DNA-templated (GO:0006355), RNA binding (GO:0003723), and RNA polymerase II transcription regulator complex (GO:0090575) were the terms with the greatest number of clusters in each of the enrichment databases searched (Kuleshov et al., 2016). Also prominent were terms related to unfolded proteins and protein turnover were response to unfolded protein (GO:0006986), regulation of protein ubiquitination (GO:0031396), chaperone cofactor-dependent protein refolding (GO:0051085), and ubiquitin protein ligase binding (GO:0031625).

**Figure 3.**
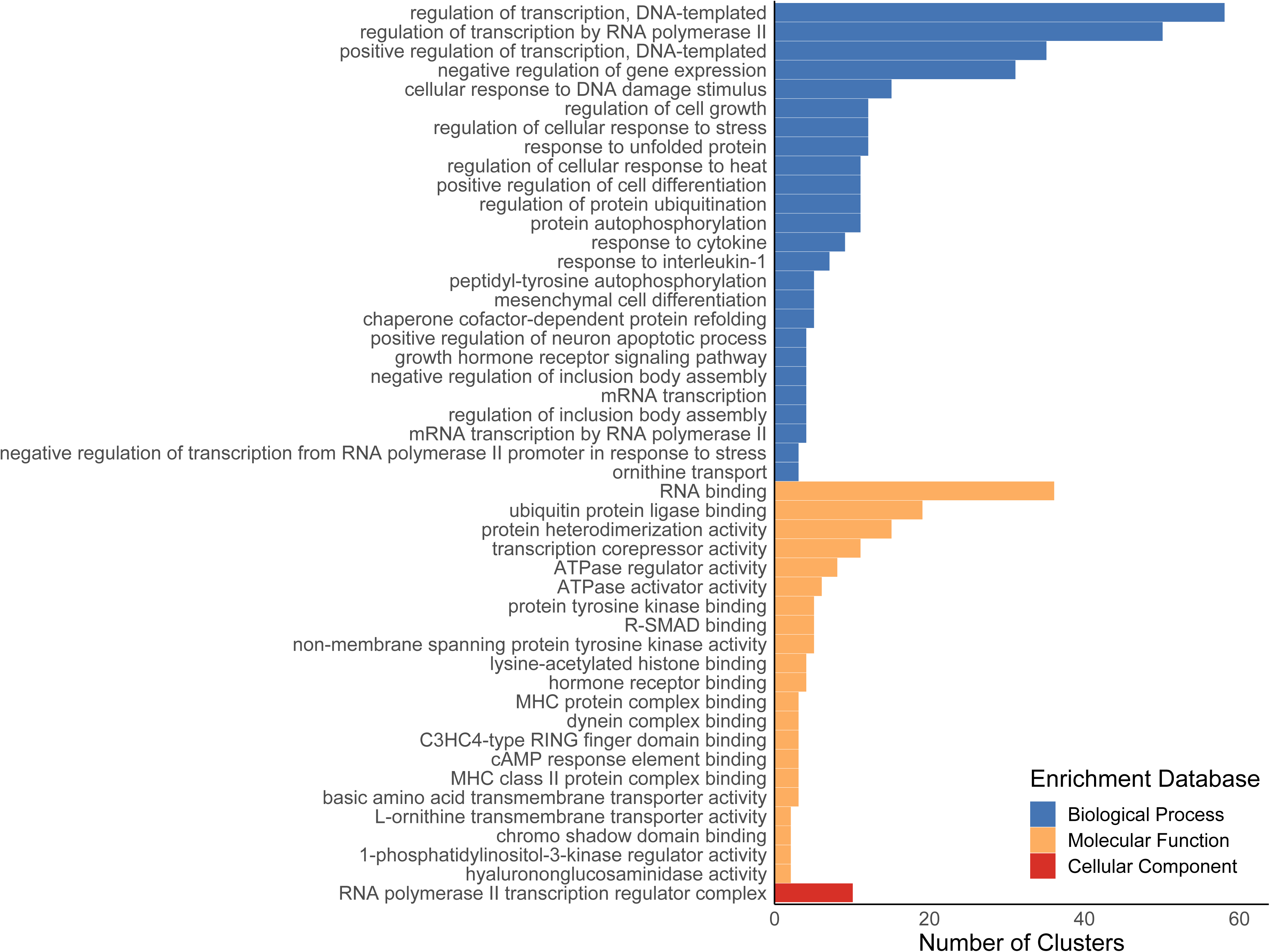
Non-redundant gene ontology (GO) terms representing clusters (∼transcripts) that showed differential expression (|log_2_-fold change| > 0) in the thermal stress treatment compared to both the handled and wild groups. Clusters were first identified as showing differential expression with edgeR, then these GO terms were called using a list of annotated genes input into enrichR. Non-redundant terms were identified with Revigo and visualized here. All terms are significant at a *q* < 0.05. Enrichment databases searched were the Biological Process 2018 (blue), Molecular Function 2018 (yellow), and Cellular Component 2018 (red). Number of clusters represents the number of genes annotated to clusters summarized within GO terms.

### Early Response Genes

Each of *jun*, *jun-B*, *jun-D*, *ier2*, *myc*, *c-Fos*, and *fosB* showed higher expression in the thermal stress treatment than in the wild group, while only *jun* showed higher expression in the thermal stress treatment compared to the handled control (Figure 4).

**Figure 4.**
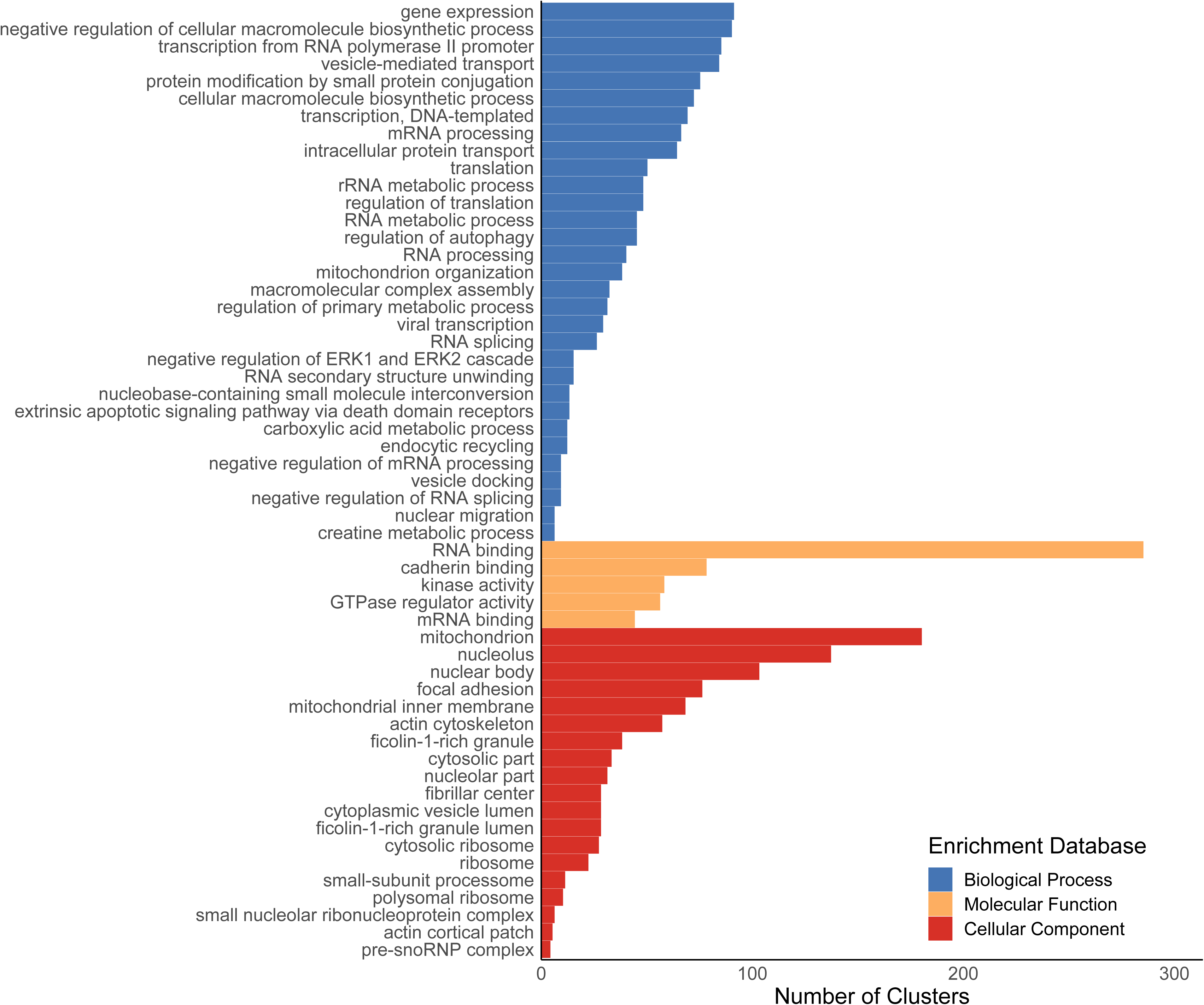
Non-redundant gene ontology (GO) terms representing exons in clusters (∼transcripts) that showed differential exon usage (|log_2_-fold change| > 0) in the thermal stress treatment compared to both the handled and wild groups. Clusters were first identified as showing differential exon usage with DEXSeq, then these GO terms were called using a list of annotated genes input into enrichR. Non-redundant terms were identified with Revigo and visualized here. All terms are significant at a *q* < 0.05. Enrichment databases searched were the Biological Process 2018 (blue), Molecular Function 2018 (yellow), and Cellular Component 2018 (red). Number of clusters represents the number of genes annotated to clusters summarized within GO terms.

### Differential Exon Usage

Among 143,841 clusters in the data, 31,042 had detectable exons, and 4,943 of these clusters (∼16%) had at least one exon that showed significant DEU between any two experimental treatments (Table 2). These clusters with significant DEU were comprised of 284,631 exons total, of which 10,314 exons showed significant DEU between any two experimental treatments. In the thermal stress experimental treatment with respect to both the handled and wild treatments, 88,031 exons in 3,230 clusters had higher expression (exon base mean 34.6 counts across samples in each exon normalized by sequencing depth, ±132.21 standard deviation) (Supplementary Table S5), while 76,307 exons in 2,530 clusters had lower expression (exon base mean 70.84 counts, ±208.98 standard deviation) (Supplementary Table S6).

**Table 2.** Summary table of pairwise results for differential exon usage among three experimental treatments, with a focus on exons unique to the CTmax treatment. Clusters (∼transcripts) were identified and quantified with Corset, and differential exon usage (DEU) was analyzed with DEXSeq. DEXSeq was used to summarize annotated clusters under different pairwise comparisons into gene ontology (GO) terms, among three databases: Biological Process 2018, Molecular Function 2018, and Cellular Component 2018. Counts of clusters associated to known genes are reported as Genes. Positive and negative expression for clusters and GO terms are with respect to both controls; positive CTmax DEU represents clusters with exons showing higher expression in the CTmax treatment compared to both controls, while negative CTmax DEU represents exons showing lower expression in the CTmax treatment compared to both controls.

|  | **Overall DEU** | **Positive CTmax DEU** | **Negative CTmax DEU** |
| --- | --- | --- | --- |
| **Number of Clusters Total** | 4943 | 3230 | 2530 |
| **Number of Exons** | 10314 | 88031 | 76307 |
| **Number of Genes** | 3136 | 2125 | 1471 |
| **Number of Biological Process GO Terms** | 56 | 46 | 1 |
| **Number of Molecular Function GO Terms** | 5 | 13 | 5 |
| **Number of Cellular Component GO Terms** | 23 | 12 | 0 |

Exons that showed higher expression in the thermal stress treatment compared to both controls were represented by 1,688 annotated genes, summarized in 72 GO terms (46 Biological Process, 13 Molecular Function, and 12 Cellular Component GO terms) (Supplementary Table S10). Using Revigo with annotated clusters containing exons showing higher thermal stress-specific expression, 25 Biological Process GO terms, 9 Molecular Function GO terms, and 11 Cellular Component GO terms were retained for visualization (Supplementary Figure S2). For exons with lower expression in the thermal stress compared to both controls, 1,170 genes were annotated, summarized by 6 GO terms (one Biological Process and five Molecular Function GO terms) (Supplementary Figure S2; Supplementary Table S11). Using Revigo, all six GO terms representing exons with lower expression in thermal stress were retained for visualization (Supplementary Figure S2). Among GO terms represented for overall thermal stress-specific differential exon usage (Figure 5), gene expression (GO:0010467), transcription from RNA polymerase II promoter (GO:0006366), and transcription, DNA templated (GO:0006351) were present. Terms directly related to metabolism were: rRNA metabolic process (GO:0016072), RNA metabolic process (GO:0016070), regulation of primary metabolic process (GO:0080090), carboxylic acid metabolic process (GO:0019752), and creatine metabolic process (GO:0006600). Mitochondria-related enrichment terms of mitochondrion organization (GO:0007005), mitochondrion (GO:0005739), and mitochondrial inner membrane (GO:0005743) were also prominent.

**Figure 5.**
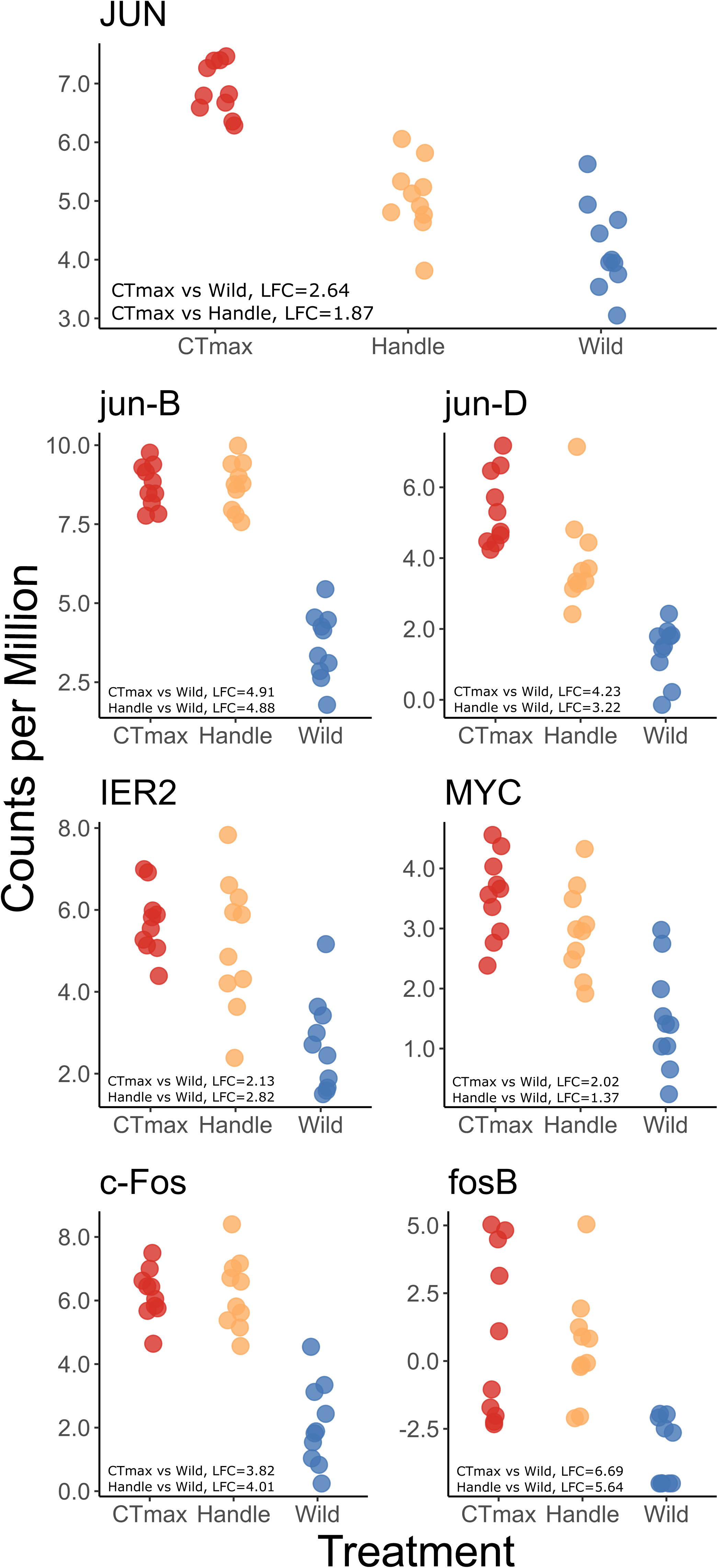
Expression of seven early response genes that are generally associated with an acute stress response. Log_2_-fold changes (LFCs) are provided for significant (*q* < 0.05) comparisons within each plot; non-significant comparisons are not shown. Individual points represent individual fish within each experimental treatment. The thermal stress treatment is abbreviated as CTmax. *JUN* is associated with *transcription factor AP-1*, *jun-B* is transcription factor *jun-B*, *jun-D* is transcription factor *jun-D*, *IER2* is *immediate early response gene 2*, *MYC* is *proto-oncogene (myc)*, *c-Fos* is *proto-oncogene c-Fos*, and *fosB* is *metallothiol transferase FosB*.

### Gene Expression-Splicing Dynamics

One splicing factor, *pre-mRNA-splicing factor 38B*, *prpf38b*, was upregulated in the thermal stress treatment compared to both the wild and handled controls (0.58 log_2_-fold change compared to wild, 0.70 log_2_-fold change compared to handled) (Figure 5). The gene *prpf38b* is associated with several genes that showed DEU between treatments: *splicing regulatory glutamine/lysine-rich protein 1* (*srek1*), *regulator of chromosome condensation* (*rcc1*), *pinin* (*pnn*), *RNA-binding protein 25* (*rbm25*), and *RNA-binding protein 39* (*rbm39*) (Figure 6; Supplementary Figure 3). The last gene, *rbm39*, is a transcriptional coactivator of *transcription factor AP-1* (*jun*), which showed higher expression in the thermal stress treatment compared to both other treatments (2.64 log_2_-fold change compared to wild, 1.87 log_2_-fold change compared to handled) (Figure 4).

**Figure 6.**
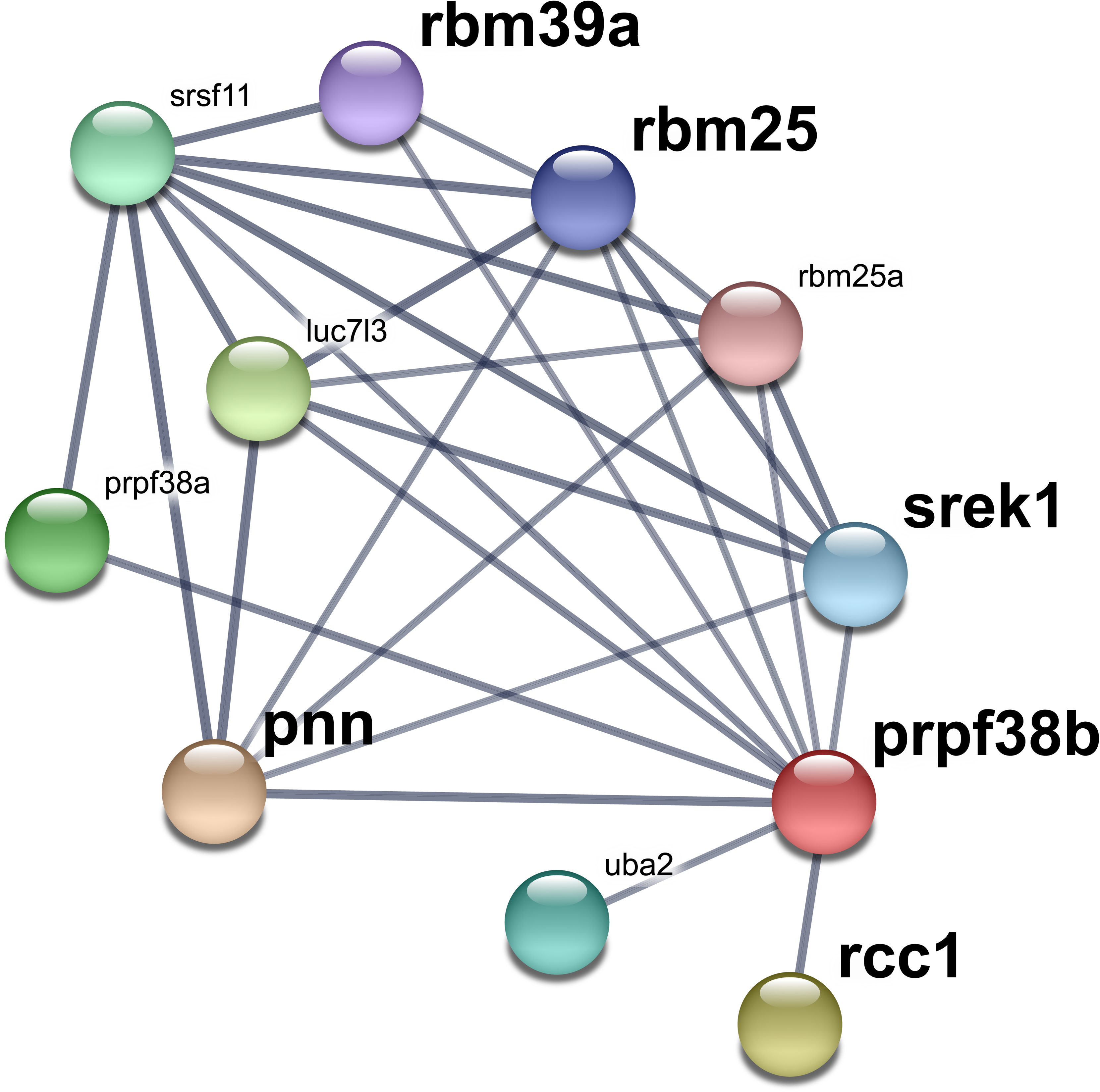
Predicted associations for *prpf38b* using the String v11.0 database. The width of lines between proteins represents confidence in the interaction, and only proteins of high confidence (>0.700) are included in this figure. The red node, prpf38b, is the query protein against the *Danio rerio* database. In bold are genes associated with *prpf38b* that showed evidence of alternative splicing *via* differential exon usage.

## DISCUSSION

Our data show that alternative splicing and gene expression may be complementary and interacting mechanisms used to mount a cellular response to thermal stress. We identified several hundred differentially transcribed genes unique to thermal stress, and these presumably represent the molecular mechanisms that redside dace use to respond to acute thermal stress. We also identified alternative splicing-based responses to thermal stress that may provide a complementary mechanism for an acute thermal stress response. Consistent with the hypothesis that differential gene expression influences differential exon usage, we identified differentially transcribed splicing factors unique to thermal stress. 1,138 clusters (∼transcripts) showed significant differential gene expression unique to the thermal stress treatment (579 positive, 559 negative). 88,031 exons in 3,230 clusters had higher expression in the thermal stress treatment compared to both other groups, while 76,307 exons in 2,530 clusters had lower expression. One splicing factor (*prpf38b*) that was upregulated in the thermal stress-challenged fish, and its increased expression was related to differential exon usage in downstream genes, representing a possible stress response pathway that incorporates both alternative splicing and gene expression.

### Gene Expression

By comparing thermal stress-challenged redside dace to handled and wild groups, we were able to identify gene expression unique to thermal stress. An observed gradient of expression responses was consistent with the handled control representing an intermediate, general stress response between the thermal stress and wild fish groups. Meanwhile, the thermal stress treatment represented a combined thermal and handling stress response while the wild treatment represented approximately baseline gene expression.

As a positive control, we used a set of seven early response genes (*jun*, *jun-B, jun-D, ier2, myc, c-Fos,* and *fosB*) that would be expected to show a stress response to verify that whole-organism acute stress was reflected in transcriptomic responses. This panel of genes was more highly expressed in the thermal stress treatment relative to the wild group. None of the seven genes in this panel showed differential expression between fish in the thermal stress treatment and handled control, indicating their role in a general stress associated with the experimental treatments as opposed to a temperature-specific stress response. Nevertheless, their higher expression in thermal stress compared to the wild group (in addition to handled compared with wild) confirms that a stress response associated with handling, transport, and confinement was reflected in gene expression. Because this panel of genes establishes that thermal and handling stressors were reflected in the transcriptomic response, thermal stress-specific genes likely represent a temperature-specific stress response when compared to both other groups.

Transcription regulation was prominent among genes differentially expressed in the thermal stress fish compared to both other groups, indicating that these genes likely play a role in coping with acute thermal challenge. While the rate-limiting step for protein synthesis is often the initiation of translation (Sonenberg and Hinnebusch, 2009; Spriggs et al., 2010), transcription regulation is another key element of the stress response (De Nadal et al., 2011). An accumulation of unfolded proteins is thought to induce a heat shock protein response (reviewed in Richter et al., 2010), and the observed enrichment terms response to unfolded protein (GO:0006986), regulation of protein ubiquitination (GO:0031396), chaperone cofactor-dependent protein refolding (GO:0051085), and ubiquitin protein ligase binding (GO:0031625) were consistent with this model. Therefore, the redside dace challenged by an acute thermal stressor exhibited a “classic” acute heat shock response as demonstrated by the multiple enrichment terms consistent with acute stress responses in the literature.

One concern with CTmax methodology is that it is based on rapid warming, which may not induce the same molecular responses that slower warming would in wild fish (Åsheim et al., 2020). However, in zebrafish (*Danio rerio*) slow warming was found to share underlying physiological mechanisms with rapid warming, evidence that CTmax induces molecular responses with consistencies across short and ecologically-relevant longer timescales (Åsheim et al., 2020). With a foundation in the conserved heat shock response among eukaryotes (Richter et al., 2010), consistency between slow and rapid warming responses in fish (Åsheim et al., 2020), and the empirical data presented in this study, the thermal stress-specific genes identified here are one mechanism of the transcriptomic response to acute thermal stress in the redside dace.

### Alternative Splicing

Given the broad importance of intron splicing in fishes and other organisms (Chaudhary et al., 2019; De Nadal et al., 2011; Healy and Schulte, 2019; Kornblihtt et al., 2013; Laloum et al., 2018; Li et al., 2020; Salisbury et al., 2021; Tan et al., 2019; Thorstensen et al., 2021; Xia et al., 2018; Zhang et al., 2019), we hypothesised that alternative splicing is an important component of the transcriptome response to thermal stress in redside dace. Therefore, we analyzed alternative splicing (measured by differential exon usage) for its possible roles in the acute stress response and interactions with gene expression. Regulation of gene expression was a prominent function among enrichment terms identified in genes showing alternative splicing in response to thermal stress. These enrichment terms are consistent with both the roles of gene expression regulation in response to stress, such as heat (De Nadal et al., 2011), and of splicing in transcription regulation more generally (Smith et al., 1989). Also prominent were metabolism-related enrichment terms among genes showing differential exon usage. Alternative splicing is one mechanism that regulates cellular metabolism, such as by splicing factors being targets of metabolic stress (Biamonti et al., 2018). Energy utilization was found to change in response to warming acclimation in fish, with decreased aerobic scope but increased energy utilization efficiency (Nyboer and Chapman, 2017; Zeng et al., 2010). Therefore, alternative splicing may represent a mechanism underlying energy use responses to environmental changes in redside dace by changing the transcribed mRNA isoforms and therefore proteomic diversity (Singh and Ahi, 2022). Consistent with this role of splicing in energy use, several mitochondria enrichment terms were significant among genes responding to thermal stress. Cellular mitochondrial content has been linked to gene expression and splicing variability (Guantes et al., 2015), and nucleus-encoded splicing machinery may splice mtRNA in humans (Herai et al., 2017). While connections between splicing, metabolism, and mitochondria are less well-characterized in fishes, these processes may play important roles in the response to increasing temperatures.

### Gene Expression & Alternative Splicing

One of our main goals was to test the hypothesis that there are direct and interacting links between patterns of alternative splicing and differential gene expression in response to thermal stress. To do this, we carefully searched for splicing factors among the genes that were found to be differentially expressed in thermally stressed fish relative to both control groups. One splicing factor, *prpf38b*, fit those criteria. Because protein abundance and mRNA levels are often correlated (Buccitelli and Selbach, 2020), and even small differences in pathway intermediates can lead to large changes in pathway flux (e.g., Hochachka and Somero, 2002), the small log_2_-fold change values we measured may be biologically important.

The splicing factor *prpf38b* may influence two important genes that are part of the thermal stress response. The gene *rbm39* was associated with *prpf38b* by co-expression in the STRING v11 database (Szklarczyk et al., 2019), and showed differential exon usage in response to thermal stress in our experiment. Furthermore, *rbm39* was differentially expressed in one of two thermally distinct populations of tambaqui (*Colossoma macropomum*) and is thought to play a role in local adaptation to thermal conditions (Fé-Gonçalves et al., 2020). In spotted seabass (*Lateolabrax maculatus*), *rbm39* was identified as a differentially expressed transcript in salt water versus fresh water (Tian et al., 2019), consistent with the differential exon usage identified in the present study. Among other roles, *rbm39* is a transcriptional coactivator for Jun/AP-1 (Jung et al., 2002). This role may be significant for the redside dace thermal stress response because *jun* was more abundant in the thermal stress treatment, relative to both other groups. Activation of c-Jun/AP-1 has been implicated in numerous, sometimes opposing context-dependent cellular stress responses (e.g., both inhibition and activation of apoptotic responses; (Leppä and Bohmann, 1999). More broadly, our data linking *prpf38b*, *rbm39*, and *jun* illustrates how the interplay between splicing and gene expression may be an essential element of the redside dace thermal stress response.

Beyond *rbm39* and *jun* specifically, *prpf38b* has been linked to the co-expression and direct regulation of numerous other genes (Ouyang et al., 2021). Therefore, while *jun* may be one regulatory element with far-reaching effects for cellular stress responses, *prpf38b* may have effects beyond *jun*, as well. As a splicing factor that was uniquely differentially transcribed in the thermally stressed group compared to all other splicing factors, *prpf38b* may be a key connection between the transcriptional mechanisms of differential gene expression and alternative splicing. In the present data, the separate gene expression and splicing analyses present enrichment term results that are presented in isolation. However, large interaction networks among genes indicate that splicing and gene expression rarely operate in isolation (e.g., Ouyang et al., 2021 for *prpf38b*; see also Boyle et al., 2017; Davidson, 2010). Therefore, further connections between the mechanisms likely exist but remain largely unexplored, possibly because of context-specificity in which splicing-gene expression interactions occur. These connections between splicing and gene expression may contribute to whole-organism stress responses, highlighting a need to study these two mechanisms in tandem.

### Conservation Implications

Understanding the mechanisms of thermal tolerance, and how these vary among populations and species, is critical for predicting the effects of environmental change and of conservation breeding, translocation, and reintroduction programs. However, these mechanisms are complex and remain poorly understood (Gangloff and Telemeco, 2018). Various studies have suggested that oxygen transport (e.g., Clark et al., 2008; Pörtner et al., 2017), coronary circulation (Ekström et al., 2019), and protein denaturation (Hofmann and Somero, 1996) set the upper thermal limits of ectotherms. Common to each of these hypothesised mechanisms is the need to mobilize energy reserves, and our data show widespread changes in patterns of both gene expression and alternative splicing related to metabolic and mitochondrial processes. This finding suggests that energy mobilization may be a fundamental factor that limits thermal tolerance. Consistent with this idea, improved nutrition has increased the thermal tolerance of redside dace (Turko et al., 2020), and several other studies have demonstrated similar patterns in other species (Hardison et al., 2021; Lee et al., 2016; Robinson et al., 2008). We speculate that there may therefore be negative consequences for the ability of fishes to cope with thermal stress in conjunction with other environmental factors that also increase energy demands (e.g., the metabolic detoxification of pollutants; Du et al., 2018). Future work using transcriptomic approaches will be useful for identifying shared pathways among these physiological processes and therefore for understanding the consequences of multiple stressors.

In addition to the shared patterns of whole organism physiology across species, cellular stress responses have deeply conserved elements across all organisms (Horne et al., 2014; Kültz and Somero, 2020). Therefore, elements of the transcriptional response to thermal stress as studied in redside dace here may be applied to understanding transcriptional mechanisms in many cyprinids and freshwater fishes. While organisms in freshwater habitats often face simultaneous stressors that limit the potential for ecological inferences in studies that use one stressor (Todgham and Stillman, 2013), rising temperatures underlie climate change’s impacts on biodiversity (Dudgeon, 2019). As such, the likely conserved elements of the redside dace’s transcriptional response to thermal stress may be representative of many cyprinids and freshwater organisms. The information presented here may be relevant to conservation of other aquatic ectothermic species.

### Conclusions

There is widespread interest in understanding patterns of inter-individual and inter-population differences for many imperilled economically and ecologically important species. For example, genetically distinct redside dace populations are known to vary in both thermal tolerance and scope for thermal acclimation (Turko et al., 2021) but the mechanisms underlying these differences are unknown. This study represents the first investigation of the redside dace transcriptome, and lays the groundwork for future inter-population studies in the redside dace and other imperilled species. Following an acute thermal stress, gene expression revealed a “classic” heat shock response, while alternative splicing revealed the potential underpinnings of changes in transcriptional regulation and cellular metabolism. Moreover, one splicing factor (*prpf38b*) was found uniquely upregulated in the thermally stressed group compared to both others here, which itself has been associated with elements of the cellular stress response (via *jun*). Alternative splicing and gene expression may thus operate in tandem in the transcriptional response to thermal stress. Therefore, the responses identified here may be among many context-dependent, biologically important interactions between alternative splicing and gene expression.

## ACKNOWLEDGEMENTS

We thank Javed Sadeghi for assistance with RNA isolation, Colby Nolan for field assistance, and Evelien de Greef for the redside dace illustration. Many analyses in this manuscript were enabled by our opportunity to use computing resources provided by WestGrid (www.westgrid.ca) and Compute Canada (www.computecanada.ca). Funding for the work was provided by the Department of Fisheries and Oceans Canada to TEP. KMJ and MJT were supported by a Natural Sciences and Engineering Research Council of Canada Discovery Grant awarded to KMJ (#05479), and AJT was supported by NSERC CREATE (ReNewZoo) and Eastburn postdoctoral fellowships.

## DATA AVAILABILITY

All scripts used for bioinformatics and analyses are available on GitHub (https://github.com/BioMatt/redside_dace_RNA). Raw sequencing reads are available on the National Center for Biotechnology Information Sequence Read Archive (accession #PRJNA692568; https://www.ncbi.nlm.nih.gov/sra/PRJNA692568).

