## Supplementary Figure for "Acute thermal stress elicits interactions between gene expression and alternative splicing in a fish of conservation concern"

**Supplementary Results**

Figure S1. Non-redundant gene ontology (GO) terms representing clusters (~transcripts) that showed higher expression (log_2_-fold change > 0) in the CTmax experimental treatment compared to both the Handle and Wild controls. Clusters were first identified as showing differential expression with edgeR, then these GO terms were called using a list of annotated genes input into enrichR. Non-redundant terms were identified with Revigo and visualized here. All terms are significant at a false discovery rate < 0.05. Enrichment databases searched were the Biological Process 2018 (blue), Molecular Function 2018 (yellow), and Cellular Component 2018 (red). Number of clusters represents the number of genes annotated to clusters summarized within GO terms.

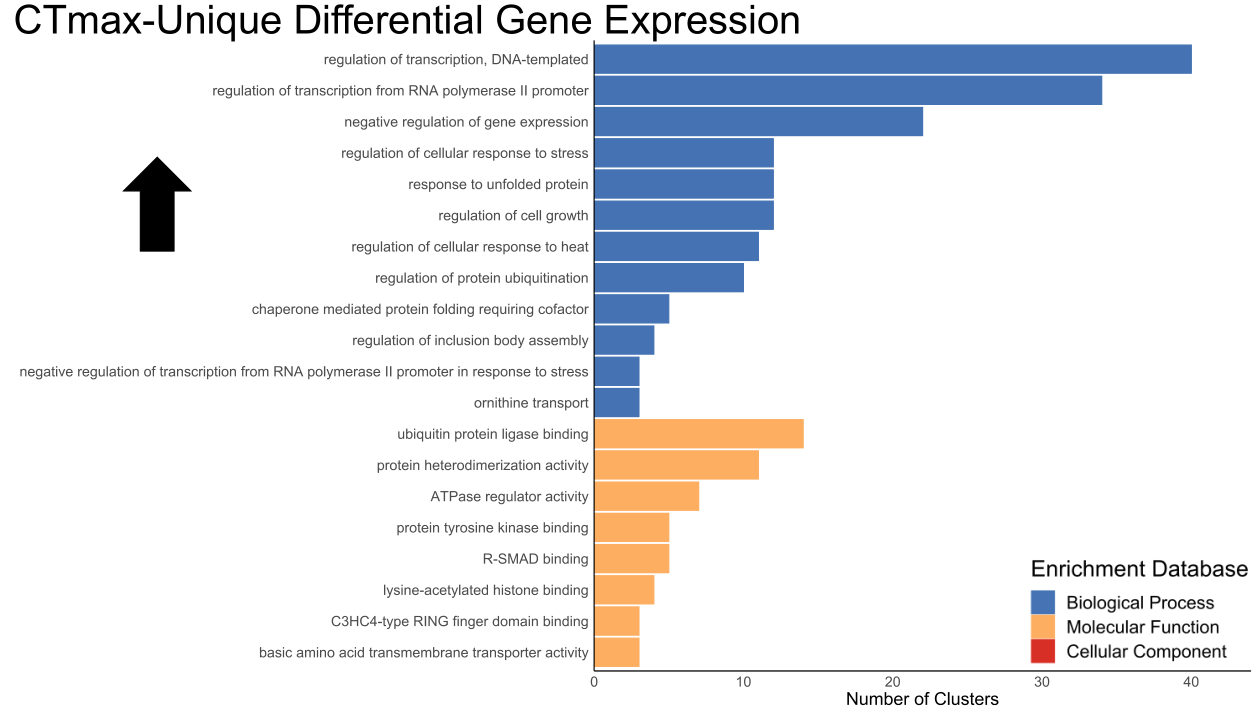

Figure S2. Non-redundant gene ontology (GO) terms representing exons in clusters (~transcripts) that showed higher (log_2_-fold change > 0) or lower (log_2_-fold change > 0) expression in the CTmax experimental treatment compared to both the Handle and Wild controls. Clusters were first identified as showing differential exon usage with DEXSeq, then these GO terms were called using a list of annotated genes input into enrichR. Non-redundant terms were identified with Revigo and visualized here. All terms are significant at a false discovery rate < 0.05. Enrichment databases searched were the Biological Process 2018 (blue), Molecular Function 2018 (yellow), and Cellular Component 2018 (red). Number of clusters represents the number of genes annotated to clusters summarized within GO terms. The black arrows represent GO terms containing exons with higher expression in the CTmax treatment (up arrow) or lower expression in the CTmax treatment (down arrow).

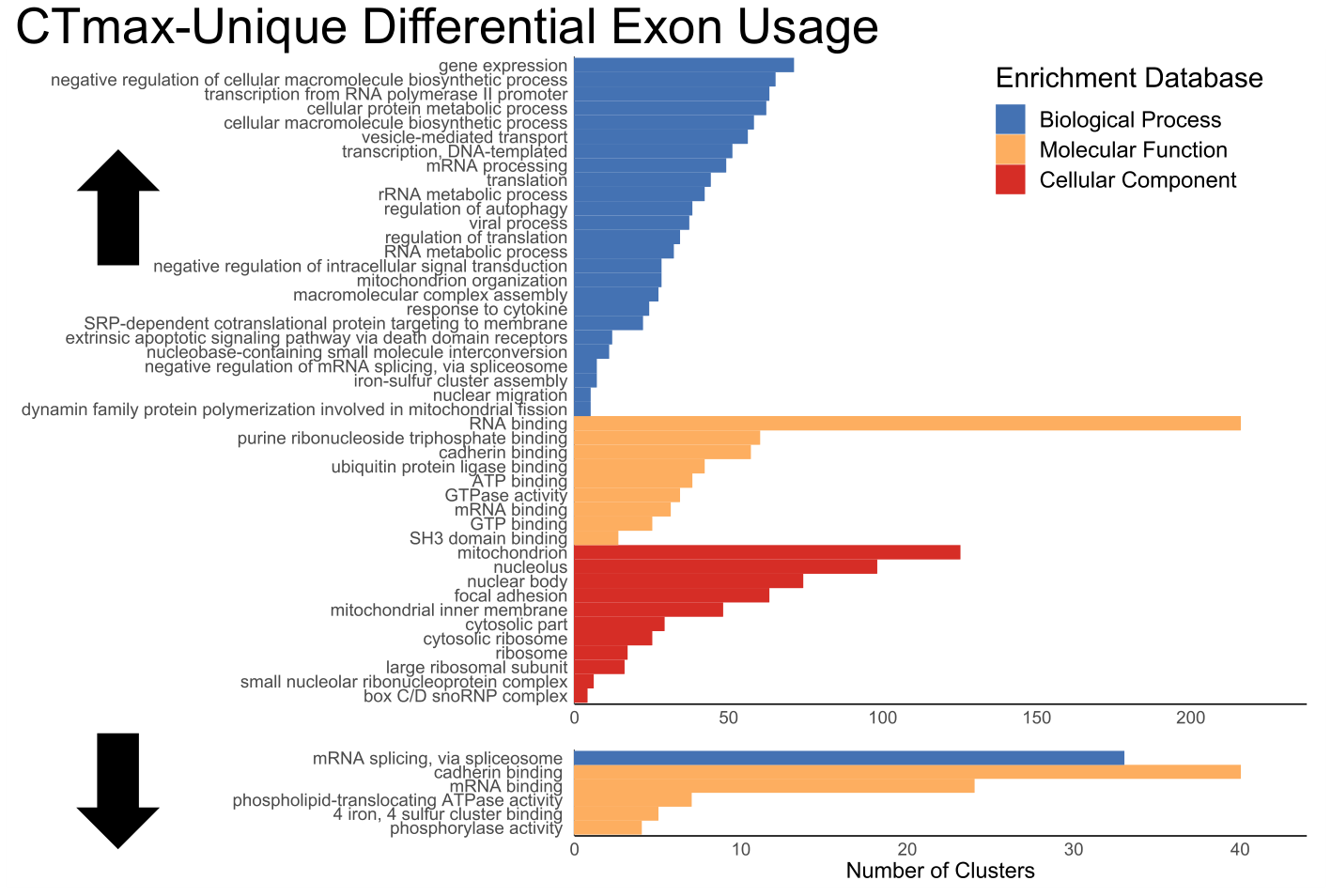

Figure S3. Counts per million for *pre-mRNA-splicing factor 38B* (*prpf38b*) and differential exon usage for *RNA-binding protein 39* (*rbm39*), *RNA-binding protein 25* (*rbm25*), *splicing regulatory glutamine/lysine-rich protein 1* (*srek1*), *pinin* (*PNN*), and *regulator of chromosome condensation* (*rcc1*). The gene *prpf38b* showed differential expression higher in the CTmax treatment compared to both controls (0.58 log_2_-fold change (LFC) higher than Wild, *q* = 1.95x10^-2^, 0.70 LFC higher than Handle, *q* = 1.71x10^-2^). Each of gene in this plot had exons with differential expression among the three experimental groups, possibly because of regulatory action by *prpf38b* (see Figure 6). Of note is *rbm39*, which acts as a transcriptional coactivator for JUN/AP-1, among other genes (see Figure 4).

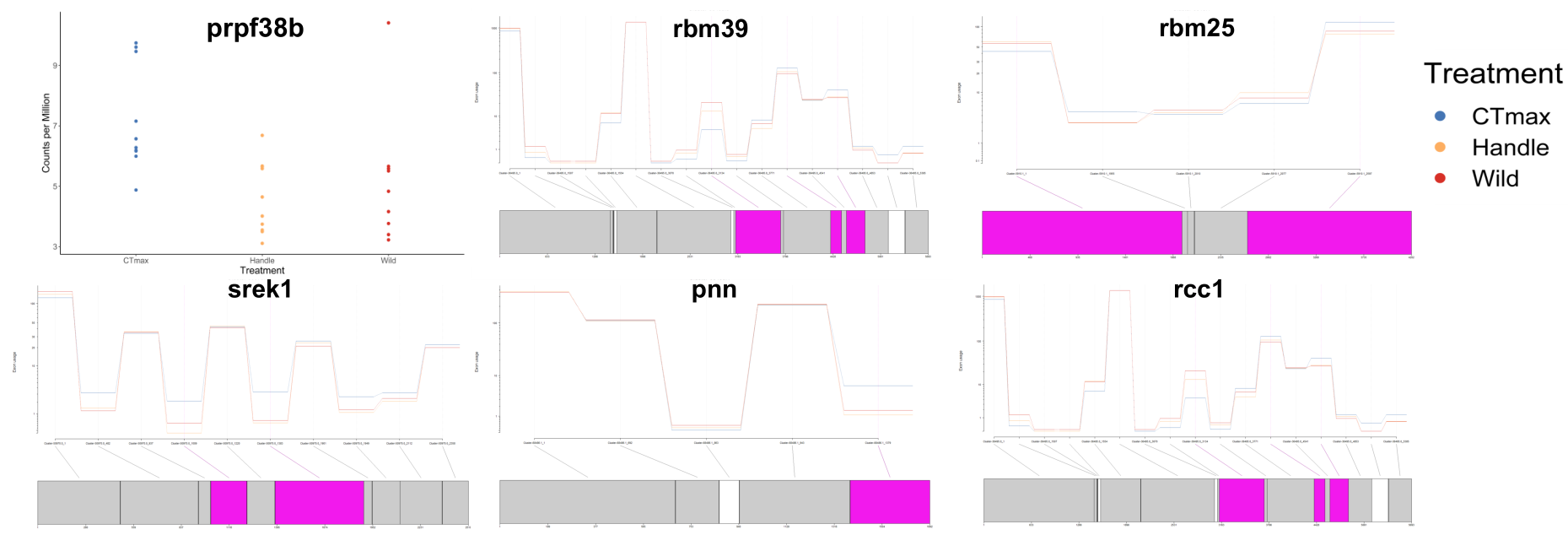
